# Lysosomal-Immune Axis Is Associated with COVID 19 Disease Severity: Insights from Patient Single Cell Data

**DOI:** 10.1101/2021.01.27.428394

**Authors:** Rahul Pande, Erin Teeple, Weixiao Huang, Katherine W. Klinger, Deepak Rajpal, Dinesh Kumar

**Affiliations:** Translational Sciences, Sanofi, Framingham, MA 01701, USA

## Abstract

SARS-COV-2 has become a leading cause of illness, hospitalizations, and deaths worldwide yet heterogeneity in disease morbidity remains a conundrum. In this study, we analyzed publicly available single-cell RNA-seq data from 75076 cells sequenced from clinically staged COVID-19 patients using a network approach and identified lysosomal-immune axis as a factor significantly associated with disease severity. Our results suggest modulation of lysosomal-immune pathways may present a novel drug-targeting strategy to attenuate SARS-Cov-2 infections.

## Main Text

Since the first cases of COVID-19 were identified in late 2019, infections have spread worldwide leading to over 100 million documented cases and more than 2.15 million deaths as of January 2020 [1]. Persistent long-term symptoms, and irreversible damage to vital organs have been increasingly reported among patients with critical as well as seemingly less severe illness [2, 3]. While several vaccines are now available, some experts have suggested that longer-term endemic levels of infection may persist, with intermittent waves of resurgence when circulating strains acquire new pathogenic mutations evading vaccine-stimulated immunity [4]. An effective COVID-19 response will therefore likely require not only widespread vaccine distribution, but also a more complete picture of SARS-Cov-2 pathogenic mechanisms in order to proactively develop and advance new treatments to address current and future challenges in terms of therapy.

Recently, single-cell RNA-seq data obtained from clinically staged COVID-19 patients was published [5]. Whereas previous analyses of this data set have focused on identification and characterization of immune cell types [5–8], we took advantage of this data to further profile COVID-19 responses and identify pathway activities associated with moderate versus severe clinical phenotypes. Data used for this analysis were single-cell RNAseq data obtained from cells in bronchoalveolar lavage fluid (BALF) samples from Healthy Controls (HC; n = 3 patients; 20991 cells) and from COVID-19 patients clinically staged as either Moderate (M; n = 3 patients; 8084 cells) or Severe (S; n = 6 patients; 46001 cells) [5]. Using R packages Seurat [9] and WGCNA [10], we performed transcriptomic profiling to identify broad cell types involved in infection response and identified gene co-expression network disease- and severity-associated modules by WGCNA (Fig.1A). Interestingly, we found that lysosomal-immune axis signaling is significantly associated with COVID-19 severity. We also identified a protein interactome for cytokine signaling associated with severe COVID-19 cases. Our results suggest distinctive transcriptomic profiles for moderate versus severe COVID-19 and that the lysosomal-immune system pathways in COVID-19 pathogenesis may offer novel drug targets.

The analysis workflow is shown in Fig. 1A. Broad cell type annotation identified numerous macrophages along with other immune cell types in all samples, with disease severity-associated macrophage populations in Severe samples (Fig. 1A). WGCNA applied to the gene-gene co-expression matrix generated from the normalized cell-gene arrays from the integrated Seurat object identified 17 discrete co-expression modules across integrated sample cells (Fig. 1A). Pearson correlation between module eigenvectors and dummy-encoded disease severity was then calculated to examine whether any of these modules might be significantly correlated with disease severity. Two of the identified modules were strongly and significantly positively correlated with Severe disease; these were the Green and Pink modules (correlation coefficient = 0.63, p = 0.00e+00 and correlation coefficient = 0.55, p = 0.00e+00, respectively). A separate Brown module identified by this analysis was significantly positively correlated with Healthy Control status (correlation coefficient = 0.71, p = 0.00e+00) and strongly negatively correlated with Severe disease status (correlation coefficient = −0.72, p = 0.00e+00).

**Fig. 1.**
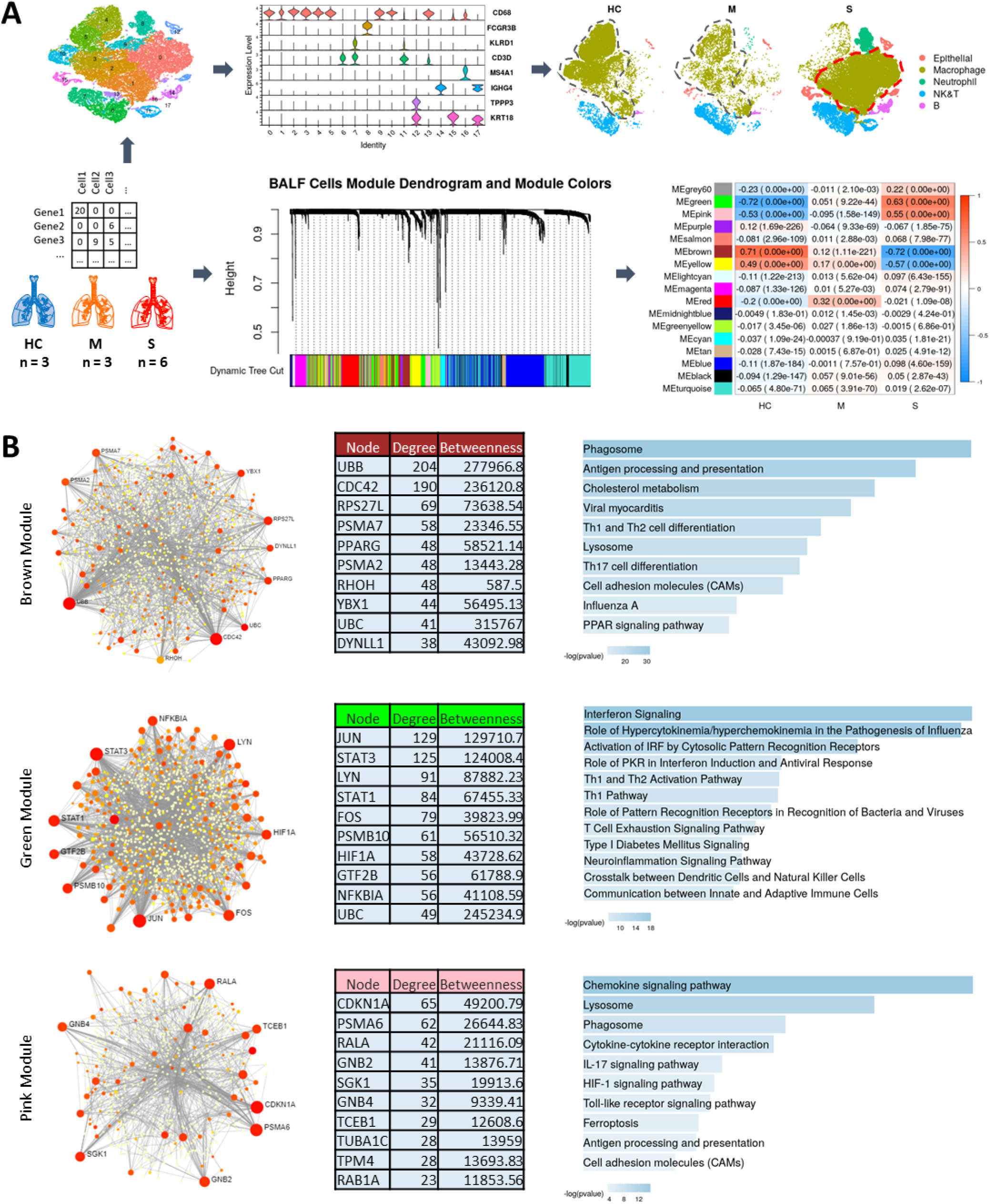
Transcriptomic Profiling and Weighted Gene Co-Expression Analysis (WGCNA). Schematic of the workflow showing single cell data from Healthy Control (HC), Moderate (M), and Severe (COVID-19) cases used for broad cell types clustering and WGCNA to identify modules significantly correlated with disease severity. Also, note the change in clustering and separation of macrophage portion in Severe cases. **(A)**. PPI networks and pathway functional enrichments for genes in highly correlated modules are shown for Brown, Green, and Pink modules **(B)**.

Positive module correlation indicates that module genes are co-expressed more actively among cells from a particular clinical group, while negative correlation suggests suppression of the module co-expression patterns. We then hypothesized that module co-expression patterns might relate to infection response mechanisms contributing to variable clinical phenotypes. To profile module functional biology, we generated protein-protein interaction (PPI) networks for module genes and performed pathway enrichment analyses for the Brown, Green, and Pink module PPI networks (Fig. 1B). PPI networks were generated for each module using the NetworkAnalyst tool [11] and pathway enrichments were queried using Ingenuity Pathway Analysis (IPA) [12] and Enrichr [13]. A number of immune system-lysosomal pathway intersections were found to be prominently associated with these COVID-19 severity-linked modules.

The Brown module, comprised of genes whose co-expression pattern is positively associated with healthy control status, is enriched for antigen presentation, phagosome formation, and cholesterol metabolism functions (Fig. 1B), with gene ontology cellular-component analysis further revealing enrichment for genes whose expression is linked with early endosomal, phagosomal and lysosomal compartments (Fig. 1B and Extended Fig. 1). Downregulated co-expression of Brown module genes in cells from severe COVID-19 patients has immune response implications. Endo-/lysosome function is necessary for antigen processing and binding to major histocompatibility complex (MHC) class II complex, and it is notable that the Brown module is also found to be enriched for MHC II antigen processing and presentation functions (Fig. 1B, Extended Fig. 1 3).

PPI network analysis for the Brown module identifies hub genes which provide further insights into the immune-lysosomal axis activity of this module (Fig. 1B). The PPI network hub protein Cell Division Control protein 42 (CDC42) is known to not only play a role in endo-/lysosomal pathway regulation but has also been shown to be used by RNA viruses for entry into the cell [14]. Inactivation of Cdc42 is necessary for depolymerization of the phagosomal cytoskeleton, subsequent phagosome maturation [14–17], and antigen acquisition and processing [18]. Another hub gene for brown module, Peroxisome Proliferator-Activated Receptor Gamma (PPARG) is a regulator of cholesterol metabolism in alveolar immune cells [19] and has been shown to regulate MHC II gene expression [20, 21]. MHC II machinery is integral for antigen presentation and is required for immunity against RNA viruses [22]. Downregulation of MHC class II machinery genes transcription is a common strategy exploited by viruses to deceive, hide from, and bypass host immune responses [23]; this same strategy is also exploited by SARS-COV2 [24]. The downregulation of Brown module genes in severe COVID-19 patients confirms via clinical evidence that these pathways are mechanisms by which SARS-Cov2 evades immune system control, as we find this downregulation to be significantly associated with severe COVID-19.

In contrast to suppression of Brown module linked pathways, the Green and Pink modules identified by WGCNA are strongly and significantly correlated with severe disease state (Fig. 1A), indicating upregulation of pathways linked with these modules among critically ill patients. The Green module eigengene has correlation 0.63 with Severe disease state (p = 0.00e+00), but negligible correlation with Moderate disease state (correlation = 0.051). Genes in the PPI network associated with the Green module are enriched for pathways involved in aberrant cytokine responses and hypercytokinemia, the so-called “cytokine storm.” These findings suggest that the Green module and its PPI network identify pathways uniquely activated in the severe disease. Abnormal cytokine release has been implicated as a main source of lung damage in severely ill COVID-19 patients [25, 26]. The PPI network for the Green module shows STAT3, STAT1, JUN and FOS as hubs (Fig. 1B). Aberrant STAT pathway activity has been shown to be central in the pathogenesis of Acute Respiratory Distress Syndrome (ARDS) and STAT3 is central to cytokine storm response in COVID-19 [27]. As an additional note, STAT3 also has been shown to associate with vacuolar H+-ATPase to regulate cytosolic and lysosomal pH [28], an activity which might further be exploited by SARS-COV-2 to deacidify the lysosome for egress of multiplying virus from the cell.

Alongside the Green module, the Pink module was also found to be positively correlated with disease severity. Genes in the Pink module are highly enriched for chemokine signaling, IL17 signaling, and lysosomal compartment (Fig. 1B, Extended Fig 1). Pink module PPI network analysis reveals hub genes for the module (Fig. 1B). Macrophages from the severe patients showed the greatest upregulation of Pink module genes. One of the hub gene, Cyclin Dependent Kinase Inhibitor 1A (CDKI1A, p21, CIP1) is a known cell cycle and apoptosis regulator. CDKI1A can be exploited by RNA viruses in macrophages to inhibit their apoptosis and turn them into reservoir for their unchecked replication without detection from host immune system [29]. CDKI1A is a key player in macrophage activation and reprogramming crucial for proper host response against viruses [30]. The enrichment of the module genes for chemokine signaling and lysosomal compartment suggests an intricate crosstalk between the two systems that has been shown to regulate the magnitude and duration of the chemokine response that is central to the ARDS or the cytokine [31, 32].

From the heatmap shown in Fig. 2B, we can see that Brown module gene expression is highest among macrophage cells in healthy controls. Green module expression is increased in severe case macrophage and neutrophil cells. Pink module gene expression is most increased among severe case macrophage cells. These module expression patterns corroborate earlier findings showing that alveolar macrophage may be the main driver of ‘cytokine storm’ in the severe patients [25, 26] and the observation suggests that macrophage specific drugs targets could be crucial for an effective therapy for COVID19.

**Fig. 2.**
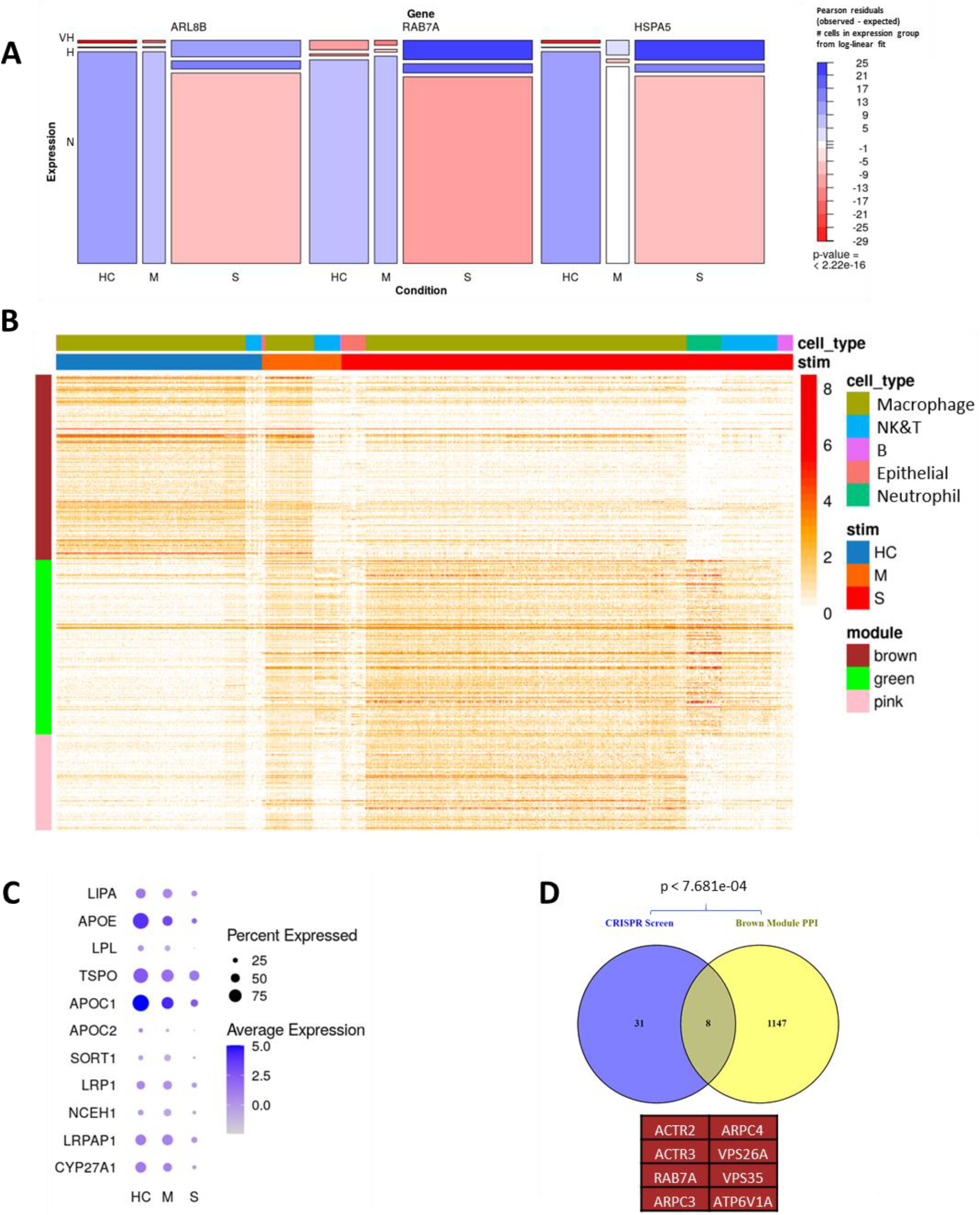
COVID-19 Severity-Associated Genes. Chi-squared testing of independence shows significant associations between COVID-19 severity and expression of ARL8B, RAB7A, and HSPA5 **(A)**. Module gene expression heatmaps for Brown, Green, and Pink modules confirm differential expression by clinical stage and cell type **(B)**. Dotplot shows expression of cholesterol metabolism genes from Brown module **(C)**. Brown module PPI network is significantly enriched for genes implicated in COVID-19 pathogenesis **(D)**.

Ghosh et al. [33] previously have shown using mouse hepatitis virus (MHV) that β-coronaviruses do not use biosynthetic secretory pathways for egress as do other enveloped viruses. Rather, β-coronaviruses hijack lysosomal machinery for this purpose: deacidifying these vessels to inactivate proteolytic enzymes and repurposing them as a novel egress passage with the help of Arf-like small GTPase Arl8b, Hspa5, and Rab7 proteins. Notably, greater expression of each of these genes ARL8B (Green Module PPI), HSPA5 (Brown Module PPI) and RAB7A (Brown Module PPI) was found to be significantly associated with disease stage among the COVID-19 cells, as can be seen by mosaic plots with chi-square test p-values in Fig. 2A. The intricate communication between endo/lysosomal and immune system is critical for proper antigen presentation and an efficient and optimum host immune response [34, 35]. MHC Class II presentation has been shown to be controlled by the lysosomal ARL8B [36] and ARL8B is likely also to be used by SARS-COV-2 for egress via lysosome, deceiving the host immune system [33].

An additional finding in the Brown module is enrichment for cholesterol metabolism pathways, which is consistent with growing evidence for participation of cholesterol in immune response pathways. Late Endosomal/Lysosomal cholesterol accumulation has been shown to be a protective mechanism among host cells for inhibiting the endosomal escape of viruses [37]. Cholesterol content is also crucial for proper antigen presentation via MHC II machinery [38]. Our findings, downregulation of several cholesterol metabolism genes in brown module in the severe patients, aligning with the findings from CRISPR screen from Daniloski et al. [39], indicate that SARS-COV2 may be benefited by the downregulated cholesterol levels in the endo/lysosomal compartment leading to an undetected and unchecked escape from the compartment evading the protective mechanism. Corroborating with the existence of mechanisms for down-regulated cholesterol in viral infection response, we found in our analysis that cholesterol metabolism genes contained in the Brown module are most heavily down-regulated in severe COVID-19 patients (Fig. 2C). Cholesterol regulation is crucial for proper antigen presentation via MHC II machinery [38], and these observed expression differences invite questions about whether and how manipulating cholesterol levels in the endo-/lysosomal compartment might offer new therapeutic strategies against COVID-19.

CRISPR screens performed by Daniloski et al. [39] to identify genes required as host factors for SARS-CoV-2 infection of human cells provide further insight into SARS-Cov-2 exploitation of the endo/lysosomal compartment. Comparison of results from our analysis with genes identified in these CRISPR screens further validates the centrality of the immune-lysosomal axis as a significant factor in COVID-19 pathogenesis. In particular, the identified CRISPR screen- genes were found to significantly overlap with the Brown module (p = 7.681e-04) (Fig. 2D). A majority of the genes identified in these CRISPR experiments, including RAB7A, are localized to the endo-/lysosomal compartment. Together, the results provide further support for endo/lysosomal compartment biology as a potential key element in the SARS-CoV-2 viral life circle.

Although the lysosome typically functions as a command-and-control center in the host immune response against viruses [40], growing evidence suggests that SARS-CoV-2 hijacks this compartment for replication, making it possible for the virus to not only evade but impair initial host immune responses [33]. Yet SARS-COV-2 also requires intact lysosomal function for its life cycle [16, 33, 40], revealing potential vulnerability along the crucial lysosomal-immune axis, as well. Our findings provide new support and clinical validation for these observations, and we identify new gene co-expression and PPI networks which may mediate disease severity. These findings have implications for the new drug development and existing drug repurposing against COVID19.

## Online content

To include any methods, source data, extended data, supplementary information, acknowledgements, peer review information; details of author contributions and competing interests; and statements of data and code availability are available

## Conflict of Interest

R.P., E.T., W.H., K.W.K., D.R., and D.K. are employee of Sanofi.

## Acknowledgements

The authors thank Dr. Srinivas Shankara for critically reading the manuscript.

## Methods

### Single Cell RNA-seq Data

Data for bronchioalveolar cells was downloaded from GEO under the accession number GSE145926. Pre-processing of data was performed as described in the source publication [5]; 12 filtered gene-barcode matrices with meta-information for each of 12 is available at https://github.com/zhangzlab/covid_balf/blob/master/meta.txt.

### Code and Pathway Enrichment Analysis Applications

Analyses and figure generation were implemented in R, version 3.6.1/RStudio.

*R packages used for analyses are freely available for download online:*

Seurat: https://satijalab.org/seurat/install.html
WGCNA: https://horvath.genetics.ucla.edu/html/CoexpressionNetwork/Rpackages/WGCNA/

*Pathway Enrichment Analysis Tools:*

NetworkAnalyst: https://www.networkanalyst.ca/ (online interface)
Enrichr: https://amp.pharm.mssm.edu/Enrichr/ (online interface)
Ingenuity Pathway Analysis (IPA): QIAGEN Inc., https://www.qiagenbioinformatics.com/products/ingenuitypathway-analysis
KEGG: https://www.genome.jp/kegg/kegg1.html (database [41])
Venny: https://bioinfogp.cnb.csic.es/tools/venny/
Project analysis code will be made available at time of publication: <GitHub link>.

### Sample Integration, Clustering, and Broad Cell Types Integration in Seurat

For quality control, cells with nFeature between 200 - 6500 and mitochondrial genes percent less than 10% were retained. Each gene-cell expression matrix was first normalized using ‘NormalizeData’ methods in Seurat v3 with default parameters, then ‘vst’ as the selection method for the FindVariableFeatures function was applied to identify the top 2000 variable genes. Samples were <u>split by disease severity and</u> integrated using the FindIntergrationAnchors function using standard Seurat workflows and 1000 anchors for canonical correlation analysis (CCA)-based integration. Cells were clustered with FindClusters function with 10 principal components and 0.6 resolution.

### Weighted Gene Co-expression Analysis (WGCNA)

WGCNA was performed in R using the ‘WGCNA’ package (v1.69) [10]. A gene-gene co-expression matrix was first generated from the normalized cell-gene arrays extracted from the Seurat data object containing all samples. Following WGCNA protocol, a scale free-topology fit index was plotted as a function of potential values for the soft thresholding power, and a soft threshold power of 6 was selected empirically for transformation of the co-expression similarity matrix into a topological overlap matrix (TOM). The TOM matrix reflects relationships of topological similarity between genes. The dissimilarity matrix (1 – TOM) is used to represent dissimilarity. This dissimilarity matrix is then used to cluster groups of genes into co-expression modules.

### Protein-Protein Interaction Network Generation

Network analysis workflow followed a protocol described in Teeple et. al. [42]. Module genes were input to the NetworkAnalyst tool using the STRING interactome database [43] with confidence score cutoff 900. which filters differential interactions by percentile from the DifferentialNet database [11].

### Pathway Enrichment Analysis

Ingenuity Pathway Analysis (IPA): Green gene set

Enrichr: For GO Molecular Functions of module gene sets

KEGG: Brown and Pink gene sets

### Statistics and Reproducibility

Data used in this study is publicly available. Statistical methods are presented here in the context of their use and interpretation

#### Hypergeometric Test for Set Overlap Enrichment

Statistical comparison of module overlaps was performed in R using the Hypergeometric test function phyper().

#### Pearson’s Chi-Square Test

For each gene, we defined three expression levels based on median (MD) and median absolute deviation (MAD). Normal level of a gene expressed in a cell is (0, MD+MAD); high level is [MD+MAD, MD+2*MAD); Very high level is (MD+2*MAD, INF). Therefore, a contingence table was able to be established for a gene with two categorical variables (Severity, Expression Level) and three factor levels each (Health Control, Moderate, Severe; Normal, High, Very High). In this case, we can determine whether Severity and Expression Level are independent for a specific gene by applying function chisq.test() in R. To better visualize a contingence table, Pearson residuals, and test result P-value, mosaic() in R package ‘vcd’ was used to make plots Figure 2A.

---

**Extended Fig. 1.**
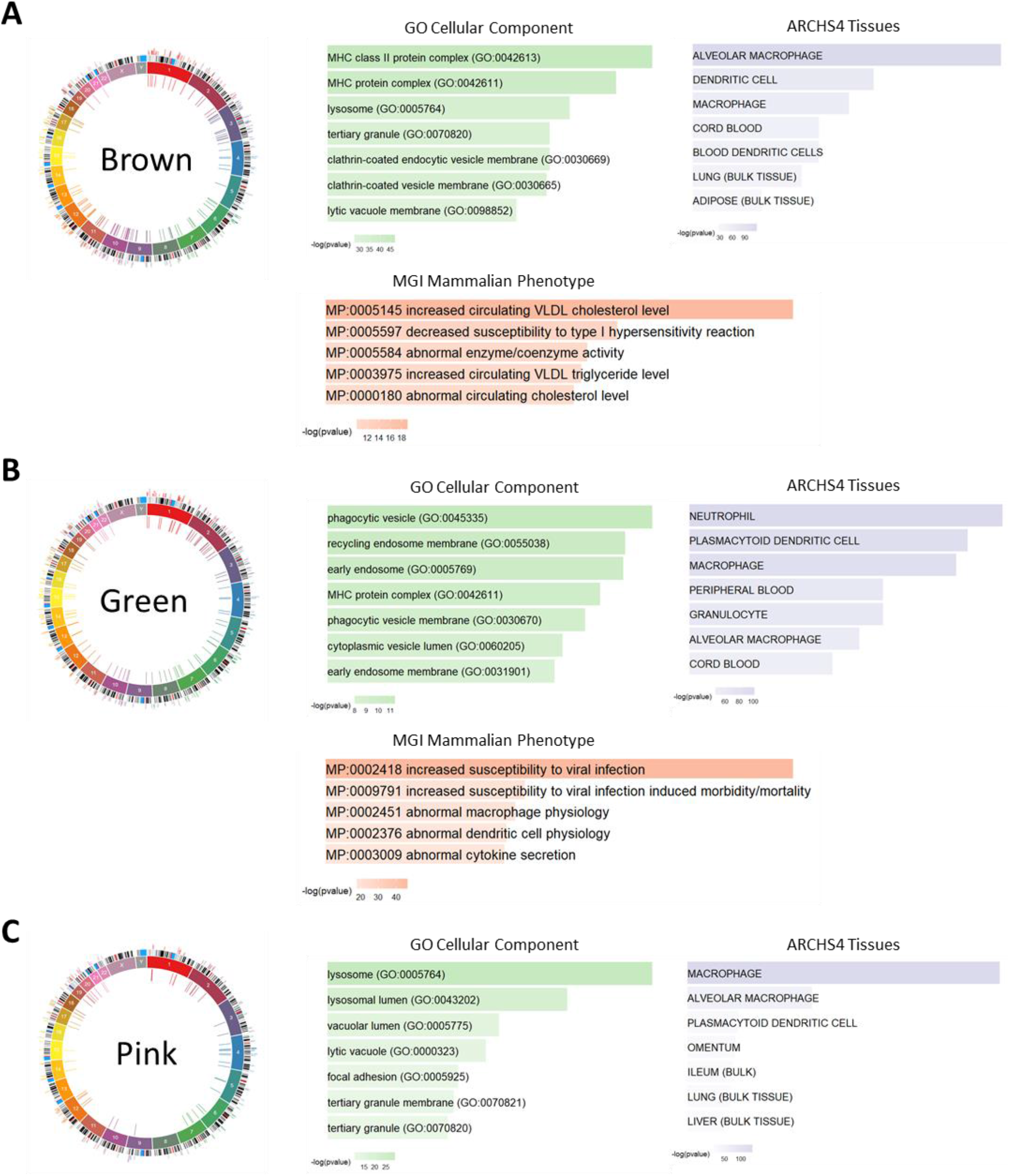
Gene Set Enrichment for the Modules. Co-Expression Modules - Brown **(A)**, Green **(B)** and Pink **(C)**-enrichment for GO Cellular Component, ARCHS4 Tissue and MGI Mammalian Phenotype provide further insight into disease mechanism and severity.

